# Effects of Spatial Heterogeneity on Transmission Potential in Vectorial-Contact Networks: A Comparison of Three *Aedes aegypti* Control Strategies

**DOI:** 10.1101/210450

**Authors:** Héctor M. Sánchez C., John M. Marshall, Sean L. Wu, Edgar E. Vallejo

## Abstract

Dengue, chikungunya and zika are all transmitted by the *Aedes aegypti* mosquito. Despite the strong influence of host spatial distribution and movement patterns on the ability of mosquito vectors to transmit pathogens, there is little understanding how these complex interactions modify the spread of disease in spatially heterogeneous populations. In light of present fears of a worldwide zika epidemic, and failures to eradicate dengue and chikungunya; there is a pressing need to get a better picture of how high-resolution details such as human movement in a small landscape, modify the patterns of transmission of these diseases and how different mosquito-control interventions could be affected by these movements.

In this work we use a computational agent-based model (ABM) to simulate mosquito-human interactions in two different levels of spatial heterogeneity, with human movement, and in the presence of three mosquito-control interventions (spatial spraying, the release of *Wolbachia*-infected mosquitoes and release of insects with dominant lethal gene). To analyse the results from each of these experiments we examined mosquito population dynamics and host to host contact networks that emerged from the distribution of consecutive bites across humans. We then compared results across experiments to understand the differential effectiveness of different interventions in both the presence and absence of spatial heterogeneities, and analysed network measures of epidemiological relevance (degree probability distributions, mean path length, network density and small-worldness).

From our experiments we conclude that spatial heterogeneity greatly influences how a pathogen may spread in a host population when mediated by a mosquito vector, and that these important heterogeneities also strongly affect effectiveness of interventions. Finally, we demonstrate that these host to host vectorial-contact networks can provide operationally important information to inform selection of optimal vector-control strategies.

**Author Summary:** Mosquito-borne diseases’ transmission patterns arise from the complex interactions between hosts and vector. Because these interactions are influenced by host and vector behaviour, spatial constraints, and other factors they are amongst the most difficult to understand. In this work, we use our computational agent-based model: SoNA3BS; to simulate two spatially different settings in the presence and absence of three different mosquito-control interventions: fogging, the release of *Wolbachia*-infected mosquitoes and the release of insects with dominant lethal gene. Throughout these simulations, we record mosquito population dynamics and mosquito bites on persons. We then compare mosquito population dynamics to the vectorial-contact networks (that emerge from subsequent mosquito bites between humans) and, after performing these comparisons, we proceeded to show that even when mosquito population sizes are almost equal in both spatial settings, the resulting vectorial-contact networks are radically different. This has profound implications in our understanding of how mosquito-borne diseases spread in human populations and is relevant to the effective use of resources allocated to stop these pathogens from causing more harm in human populations.

## Introduction

*Aedes aegypti* mosquitoes are responsible for the transmission of some of the most epidemiologically important vector-borne diseases in recent years: dengue, chikungunya and zika. In the past 50 years, the incidence of dengue has increased drastically [1, 2]. Recent estimates suggest around 390 million annual cases globally and rising; with nearly 96 million of these exhibiting clinical symptoms [3]. Similarly, chikungunya and zika are emerging diseases that are rapidly disseminating in Latin American countries, raising health concerns and placing a heavy burden upon their national health institutes [4–6].

These diseases are transmitted as a side-effect of *Aedes*’ need to obtain blood to complete its life cycle (gravid female mosquitoes require blood from a host for their eggs to be viable). Arboviruses, in turn, use these mosquitoes as vehicles to develop and travel between human hosts. Unfortunately, to date, there exists no effective vaccine to block transmission of any of these pathogens (Dengvaxia, the only available vaccine against dengue has been the subject of recent health concerns [7, 8]); so the control effort has largely focused on the disruption of the mosquito life cycle. This has proven to be a difficult endeavour.

Several mosquito-control interventions exist to date and more are being developed every year. Amongst the traditional *Aedes*-control interventions, spatial insecticide spraying (also known as fogging) is one of the oldest. Sadly, in spite of its long history and widespread use, the efficacy of fogging campaigns has been generally been limited [9, 10]. This, coupled with other traditional approaches’ limitations, has created a pressing need for novel approaches to contain the pathogens’ spread. In recent years, two of the most promising novel interventions have been: release of *Wolbachia*-infected mosquitoes and release of insects carrying a dominant lethal gene (RIDL). *Wolbachia*-based strategies work by infecting mosquitoes with a bacteria which has been shown to limit the potential of some arboviruses (such as dengue and chikungunya) to develop and subsequently be transmitted [11]. Female-Specific RIDL techniques, on the other hand, work by genetically modifying mosquitoes so that females carrying the dominant gene do not develop viable wings upon progression to adult stages [12].

Despite the fact that these novel interventions have shown promising results in field trials [13, 14]; evaluating their cost and effectiveness in a wide variety of different settings is both crucial and difficult. Evaluating interventions in the field is expensive both in time and economic resources. This, paired with the fact that mosquito-borne diseases usually affect low-income countries, makes it paramount to predict their impact before actually using them in practice.

Mathematical and computational models are effective tools to predict the impact of interventions on pathogen transmission. Classical models of pathogen transmission based on systems of ordinary differential equations (ODEs) have been standard tools for these types of analysis. Despite this, spatial heterogeneity in host-vector interaction, noted to be fundamental to patterns of pathogen spread, is difficult to incorporate in them [15–17]. Agent-based models (ABMs), on the other hand, can handle this information in a natural way so they have proven to be useful for these analyses. Along these lines, we have developed SoNA3BS: Social Network *Aedes aegypti* Agent-Based Simulation (SoNA3BS was coded in *NetLogo* and its source code will be freely available upon publication in our git repository: https://github.com/Chipdelmal/SoNA3BS). Our model simulates interactions between humans and mosquito agents on a defined landscape. With it, we are able to track the bites females take on humans; which allows us to reconstruct the vectorial-contact networks and to borrow tools from graph theory to perform structural analyses. This, to the best of our knowledge, is on the vanguard of epidemiological analysis of vector-borne diseases (although it has successfully been used before for direct-contact pathogen transmission [18–22]).

The importance of spatial heterogeneities on pathogen transmission [17, 23, 24], has motivated our use of SoNA3BS to explore the effect of spatial arrangement on vector-borne disease transmission. Specifically, we used SoNA3BS to test the hypothesis that spatial distribution of human houses and mosquitoes breeding sites has a significant effect on reshaping the way human contacts occurred in both, in absence and in presence of three mosquito-control measures: spatial fogging, *Wolbachia*-releases and RIDL. To test these effects, we first simulated two different spatial scenarios: a homogeneous one (in which every household and human is placed in the exact same place) and a heterogeneous one (with a more natural spatial distribution obtained from a real human settlement). We then obtained both the population dynamics and the vectorial-contact networks that result from applying each of the interventions in the environment. This information allowed us to show what is, in our opinion, the main contribution of this work: despite the fact that the mosquito population dynamics remain almost identical in both situations on all cases, the networks that arise from them have different structural properties. This, in turn, shows that mosquito biting heterogeneities can arise solely from spatial distribution and highlights the importance of taking into account spatial information in the planning of mosquito-control interventions’ deployment on the field.

## Materials and Methods

To investigate how spatial distribution of hosts and sites affects the ability of vector control interventions to disrupt pathogen transmission, we simulated two different scenarios under the presence of *Aedes* control campaigns. After doing so we analysed both the population dynamics and vectorial-contact network structures.

In the following section we will discuss how these spatial settings were selected and implemented on the simulation; along with the measurements that were performed as part of the analysis.

### Simulated Scenarios Settings

We first describe how sites were defined in each scenario. We also describe the behaviour rules used to simulate mosquito and human activity.

### Landscape and Humans

We generated a simulated version of a location near the Mexican town of Catemaco, Veracruz by obtaining approximate positions of houses’ locations using *Google Maps*(geographical coordinates: 18°25′52.0”*N* 95°05′25.1”*W*). We chose this area because it is a small-sized region in M´exico in which the *Ae. aegypti* presence is widespread [25].

A population of 30 humans was distributed amongst 12 houses proportionally to the household area that was detected (viewed from the satellite image). This gives us an average value of 2.5 persons per house. For each household we assumed one viable egg-laying site in its vicinity, as most of the populations with no piped water supply use containers to provide for their needs [26]. Under these conditions, we simulated two different scenarios:

- Homogeneous (HOM): Every house and person was placed at the center of the environment as depicted in figure 1a. Humans remained static while mosquitoes were allowed to move to fulfil their biting, sugar feeding and eggs-laying needs.
- Heterogeneous (HET): Houses in the environment were placed as shown in figure 1b. A maximum of two humans per household were allowed to visit other households (with a probability of 10% per day), while the remaining humans stay at home. This house was chosen randomly from the pool of houses in the simulation each time a visiting event was triggered.

**Fig 1.**
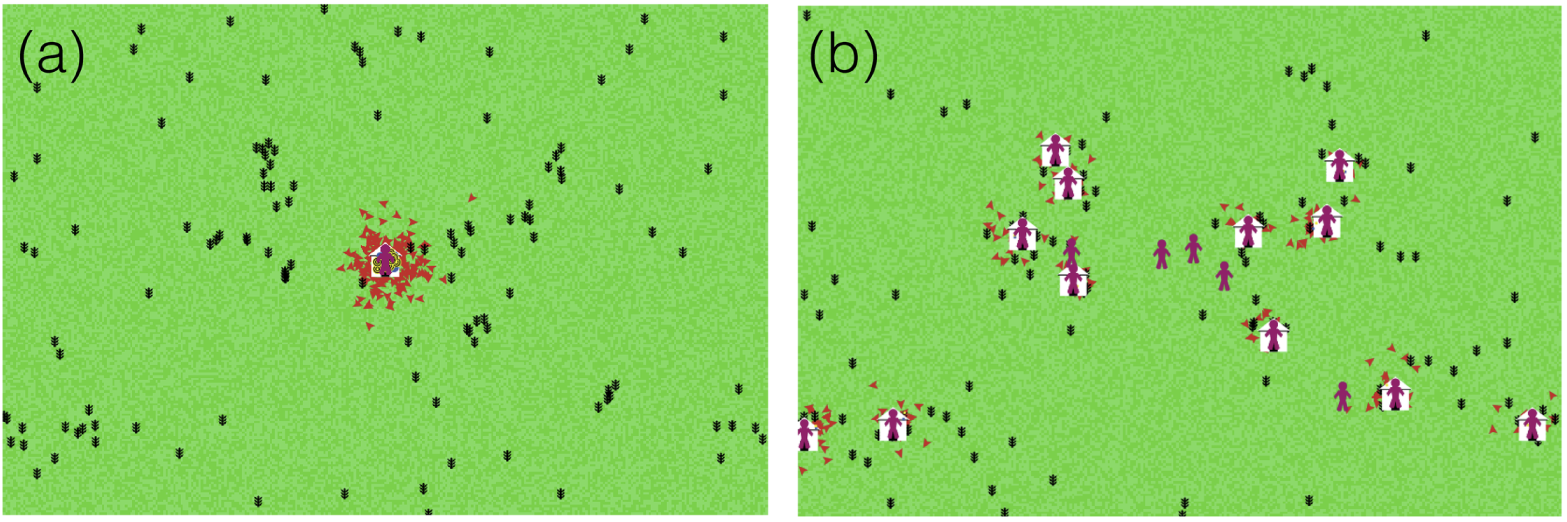
Simulated spatial scenarios. SoNA3BS screenshots representing the two modelled layouts: homogeneous (a) and heterogeneous (b). Humans are shown in purple, houses in white, sugar sources in black and mosquitoes in red (triangles). Breeding and mating sites were scaled down but are always near the location of the houses according to the methodology described while the other agents and landscape elements were scaled up for readability purposes.

### Time and Temperature

Computational agent-based models do not run in continuous time as ODEs do, so special care has to be taken in selecting a time-step (or “tick”) resolution. This is because if the amount of time per tick is set too high all the fine detail that could be captured in the model is lost; whilst if a very high time resolution is selected the computational burden of the model outweighs the benefits of high temporal resolution. In our simulation we gravitated towards a high time resolution value of 5 minutes/tick (other similar ABMs generally step sizes of hours or days [25, 27, 28]). This is important for our analysis because we are interested in the analysis of fine-detail interactions between individuals so we need to be able to simulate them as precisely as computationally possible.

The simulated timespan is also important in agent-based models as it can affect the analysis of the dynamics of the system (enough time must be given for transitory dynamics to settle). In our study we needed variations in the populations’ sizes and, as a consequence, mosquito bites to achieve stability while still maintaining a low computational cost. To achieve this, each run comprised a period of 360 days with a burnin period of 100 days. Both of these periods were defined empirically after running extensive preliminary tests on the simulation. With the timing information of our experiments in place we moved on to defining its weather characteristics. Temperature is an important part of mosquitoes metabolic development (in warmer conditions, mosquitoes develop faster [29]) and our simulation does incorporate these effects on mosquitoes biology. For the purposes of the experiments presented in this work, we fixed the temperature’s value to 25°C (close to the average temperature of the region which was calculated with data obtained from: http://clicom-mex.cicese.mx). We made this decision because we wanted to limit the effect of variables other than spatial distribution on the results (having a realistic seasonal pattern would have not only affected the mosquito population dynamics but it would have also added interactions with the timing of mosquito-control interventions [13, 14, 30]). Along the same lines we assumed that humidity and rainfall were constant and adequate to maintain *Ae. aegypti* populations throughout the year.

### Mosquitoes

Simulating a realistic population of mosquitoes was a crucial part of our experiments and as such, special care had to be taken in defining their quantities and their behaviours.

Under the described conditions, our IBM produced a baseline population at equilibrium of 30 adult mosquitoes per aquatic habitat. This number is close to densities observed in Cayman Islands (where RIDL field tests took place and with similar weather patterns to the ones found on Catemaco [14]). It is important to note, though, that this carrying capacity value is not a hard threshold in our IBM, but an emergent property that arises from interactions between mosquitoes and the environment as a whole.

In terms of the mosquitoes behaviour, our simulation treats mosquitoes going through their life stages as finite state machines; each having some associated set of behaviours. Both male and female mosquitoes go through three aquatic stages: egg, larva and pupa; in which they spend their time maturating according to their metabolic rules of development [29]. After emerging from their aquatic phases, males spend one day waiting for sexual maturity and then they spend their lives in feeding and mating bouts. Females, on the other hand, emerge sexually mature and mate just once throughout their lifetimes. After doing so they spend the rest of their lives in sequential cycles of blood-feeding, resting and egg-laying; just pausing to sugar feed when needed. These behaviours are summarised in figure 2. All adults are more active during the daytime than at night, which corresponds to *Aedes aegypti* behaviours in real life [31].

**Fig 2.**
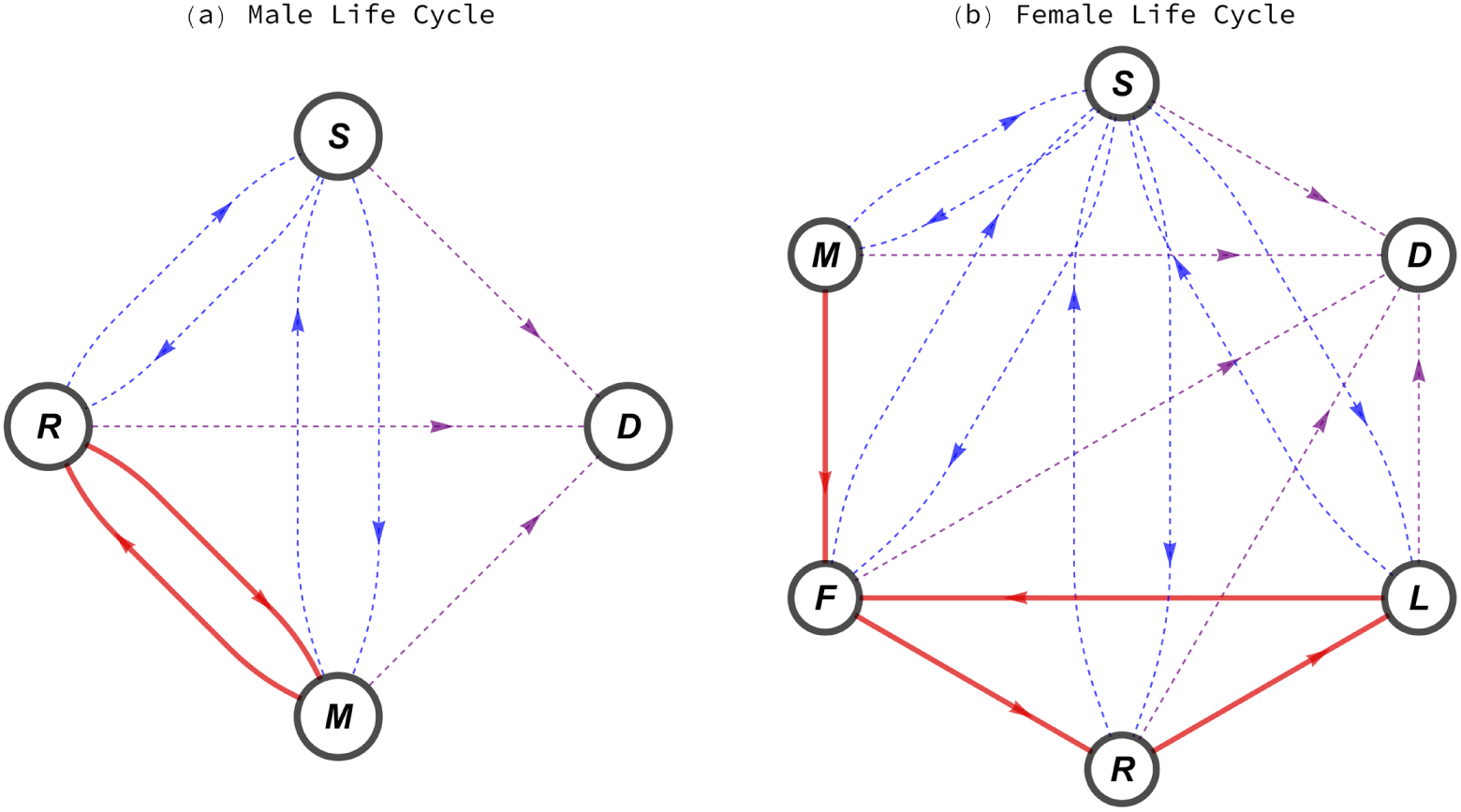
Mosquitoes life-cycle transitions diagrams. Male cycle (a) is comprised of repeated mating (M) and sugar feeding (S) cycles after their initial rest bout (R). Female cycle (b) involves iterations of blood-feeding (F), resting and egg laying (L) actions after they have mated. Both males and females can go into a sugar feeding bout if needed or die (D) due to biological or control driven phenomena.

Biological parameters such as metabolic rate, flight range, eggs laid per gonotrophic cycle, flight speed, death probabilities and population densities; were calibrated specifically for *Ae. aegypti* mosquitoes (these parameters are available as part of the supplementary material). The specific values of these variables will be accessible in the setup routines of the simulation upon publication. There, they can be viewed and modified to accommodate reproducibility and model extension purposes.

### Simulated Mosquito-Controlled Interventions

Once the landscape and behaviour of the agents was defined we focused on the way the vector-control interventions affected mosquito individuals. Three different mosquito interventions were simulated along with a baseline scenario. Their effects on mosquitoes were defined as follows:

- Baseline (Base): Mosquito dynamics and death probabilities remained unchanged.
- *Wolbachia* (Wolb): The bacteria was inherited between generations according to 1 cytoplasmic incompatibility’s rules of inheritance [10]. Adult lifespan was halved on average and the ability of *Wolbachia* to block the disease’s transmission was assumed to be 90% (although studies suggest that this percentage could be higher depending on the strain used [13, 32]).
- fsRIDL (RIDL): Mosquitoes carrying a lethal gene allele were allowed to mate and reproduce but their female offspring did not develop viable wings [14]. Only males could mate further and propagate their genes. These RIDL genes were transmitted to the offspring in accordance to Mendelian inheritance laws.
- Spatial Spraying (Fog): An instant-action layer of insecticide is applied to the whole environment (the WHO recommends the application of insecticide in an radius of 400m around the houses where Dengue is detected [33]; and the dimensions of the simulated environment are 1000m x 1500m so this approximation is within this recommended range). This insecticide has a fast exponential decay (starting with a death probability of 6.25% immediately after application and a half-life of 120 minutes); and it is assumed to affect only adult mosquitoes. These parameters were defined empirically but could be fitted to any specific type of insecticide if data became available.

Control measures were applied uniformly in the environment. *Wolbachia* and RIDL had a fixed number of mosquitoes released uniformly over the landscape; while fogging was assumed to work with equal efficacy across the whole landscape. This decision was taken to focus on how the spatial distribution of individuals affects the effectiveness of the control campaigns, not the specific way in which interventions are applied in the landscape (which is known to be important [34, 35]).

In terms of time and density of the releases, intervention events took place on a weekly basis to match field campaigns; and the number of RIDL and *Wolbachia*-infected released individuals were also scaled to match field tests (225 individuals per release [13, 14]). The main difference between the way the interventions campaigns were simulated was that *Wolbachia*-infected mosquitoes were released for five weeks (in this period the pathogen achieved fixation in all situations), while RIDL and spatial spraying were applied during the rest of the simulated time to make a fair comparison between the interventions (*Wolbachia* gets fixated in the population and continues to be propagated while RIDL is self-regulating, and fogging stops working almost as soon as campaigns finish).

Each combination of spatial scenario and control measure was repeated 30 times to reduce the variance on the analysis.

### Comparison Metrics

With the simulation’s settings defined we now describe the analysis we performed on the obtained data. As discussed earlier, we performed a contrast analysis to compare population dynamics and vectorial contact networks across different scenarios. We think that making these comparisons is a meaningful way to separate the effects of spatial location from simple population counts. This is because in a scenario in which the spatial distribution of individuals had little to no effect, we would expect the quantity of mosquitoes and the biting networks structures to change proportionally to each other. However, if the spatial effects are meaningful, some independence in the way the metrics behave is expected.

#### Population Dynamics

To analyse the impact of the control measures on mosquito population sizes, we stored their demographics twice a day. We focused on the analysis of adult mosquitoes which were broken down into the following categories: total adults, adult females, adult males, RIDL-infected and *Wolbachia*-infected. It should be noted, though, that we did store information on other life stages (eggs, larvae and pupae) in case further analysis is deemed useful. All of these data will be available upon publication.

#### Networks

The vectorial-contact networks are, in our opinion, the most novel part of our analysis. Despite the fact that network epidemiology has become more widespread in direct-contact diseases [36–38]; in vector-borne scenarios performing this kind of transmission analysis is difficult. The use of an IBM allows us to track the biting history of each mosquito, so we can recreate not only the epidemiological transmission network but also the network that arises purely from mosquito bites (of which the epidemiological one is a sub-network). Networks were obtained according to the following procedure:

1. Mosquito bites were recorded along with the time in which they occurred.
2. If a person was bitten after another person was also bitten by the same mosquito (in a previous gonotrophic cycle) a vectorial transition was created between them (in the case of *Wolbachia*-infected mosquitoes we are assuming pathogens transmission reduction, so 90% of the bites from them were discarded [13, 32], as we are interested in the potentially-infective ones). If the same person gets bitten in subsequent gonotrophic cycles these links are discarded as these bites are not epidemiologically relevant (self-loops would not disperse the disease in the population).
3. The resulting network is created by translating vectorial transitions into edges between persons (which are the vertices).

Once these graphs were generated for each scenario we calculated a collection of network measures that are related to disease transmission within a population [19, 39–41]. We did this for both the weighted networks (with the weight being the number of transitional bites between two given individuals) and the binary networks (one or more transitional bites are treated simply as one transition).

The first metric we evaluated was the in-degree distribution of the persons (number of incoming bites after the first mosquito’s gonotrophic cycle). This is related to the general risk of a person to contract a vector-borne disease and is of upmost importance to understand diseases transmission. We defined the in-degree as the total number of incoming bites for each person as we are working with weighted networks (also known as multigraphs).

In addition, we also analysed the following network measures:

- Graph Density: Represents the ratio between the number of edges of our graph and the number of edges of a fully connected graph with the same number of vertices. This measure indicates how our network compares to the absolute worst case scenario where the possibility for transmission between every human in the population exists.
- Mean Path Length: Average length of the shortest paths in the network. It is related to the speed and reach a disease could have in the population because it represents to how many “jumps” a pathogen would have to make to cross from one human host to another.
- Small World Coefficient (SW): Measures to what extent nodes are neighbours of one another in relation to how long are their paths to other nodes in the network. When this quantity is high, networks have the characteristic of having nodes that are separated by a low number of steps but that are not necessarily neighbours with each other. The small-world effect is of epidemiological concern because it allows a pathogen to spread on a network even when connections between individuals are sparse (the mean path length grows logarithmically with the number of nodes) [39].

As a final step of analysis of the relation between spatial distribution and the frequency of transitional bites we used spectral clustering on the networks [42]. This allowed us to identify patterns that were arising on the biting behaviour of the mosquitoes and it is important because, if no clustering patterns could be found on the heterogeneous layouts, then we could conclude that there was no direct relation between person’s location and the transitions amongst them.

All of these networks analyses were performed on *Mathematica 11* using the functions provided with the software and extending its capabilities by using *PajaroLoco* (H´ector M. S´anchez C. [43]), a package developed for networks’ structural analyses.

## Results

In the following section we will show the results obtained from the interactions of mosquito and human agents in our simulation. First we will show how population dynamics behaved in the presence and absence of spatial heterogeneity. Then we will make the same comparison on the vectorial contact networks. After making these contrasts we will describe briefly the differential effect of each intervention in terms of population sizes and efficacy on the disruption of vectorial contact networks.

### Population Dynamics

Most of the interventions behaved similarly in terms of the way they affected mosquito population sizes across their homogeneous and heterogeneous cases, falling in line with our initial hypothesis that they should be similar given the conditions of the experiments. The one notable exception was RIDL. We can observe in figures 3a and 3c a slight difference on the long term effect of the releases. In the long run, the heterogeneous scenario suggests that RIDL alleles had a harder time getting transmitted in the mosquitoes population; but presenting firm conclusions on RIDL dynamics requires longer simulation times (an objective for future research). For the purposes of this particular experiment we can say that, at least in the simulated timespan, the population sizes in both RIDL settings were very much alike (something that will be discussed looking at the mean population sizes of the experiments).

**Fig 3.**
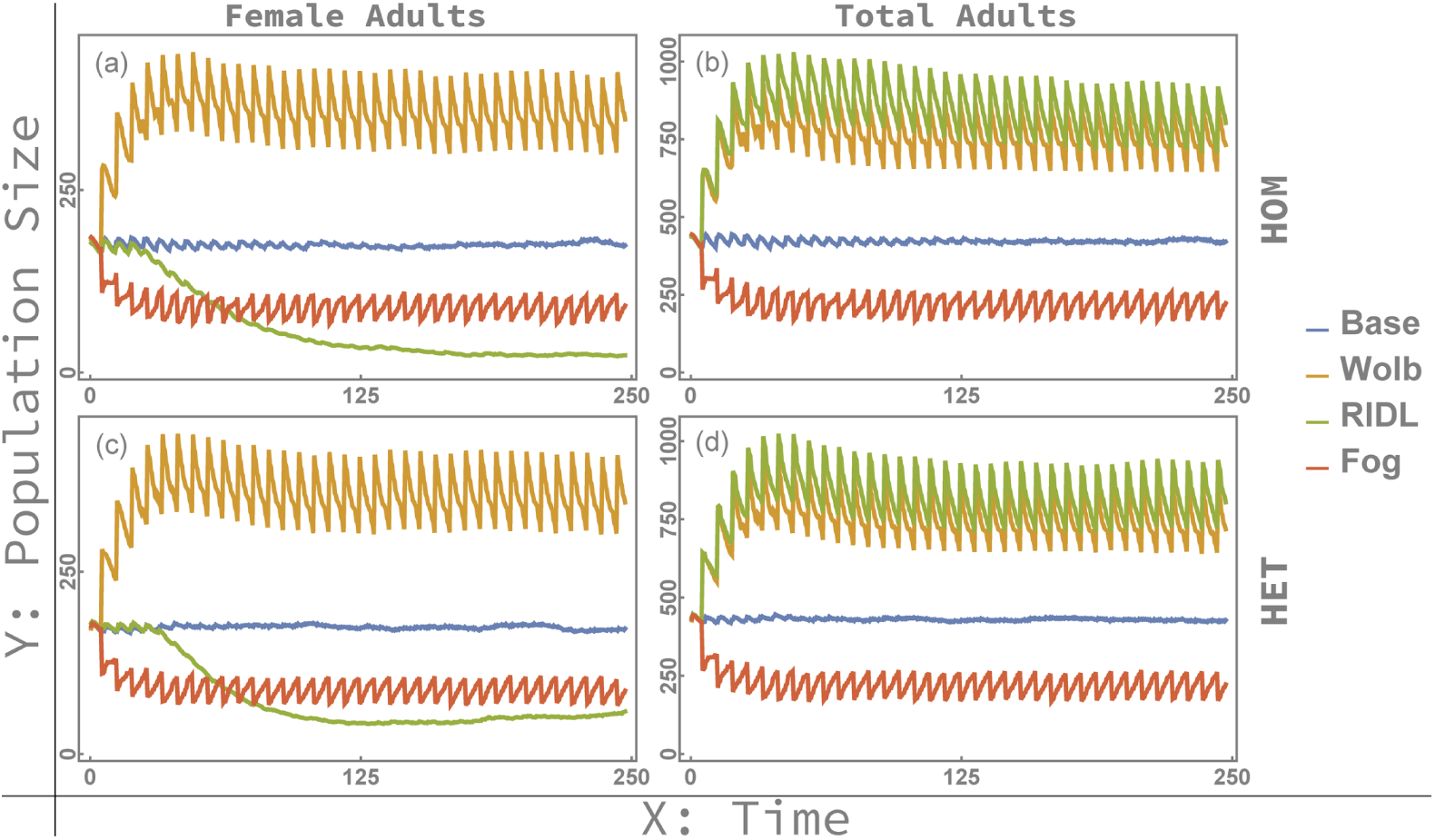
Population dynamics of adult mosquitoes. Figures (a) and (b) show the obtained populations in the homogeneous cases while figures (c) and (d) the heterogeneous ones. Given the experimental settings, the differences between the two spatial settings are barely noticeable through visual inspection.

Besides RIDL’s specific case, visual inspection of the plotted data was not enough to establish any significant differences on population sizes between spatial layouts, so we calculated the average population size from each experiment (area under the curve divided by the number of time points). The results of these calculations are shown in figure 4, confirming that no meaningful deviations on population sizes were observed between spatial settings. This lack of differences is most likely due to the fact that the tested environment was relatively small, interventions were applied uniformly on sites, and there was little environmental pressure on the mosquitoes (abundance of sugar food sources and human hosts). We would expect the dynamics to change if either environmental or behavioural variables become a significant external stressor on mosquito survivability.

**Fig 4.**
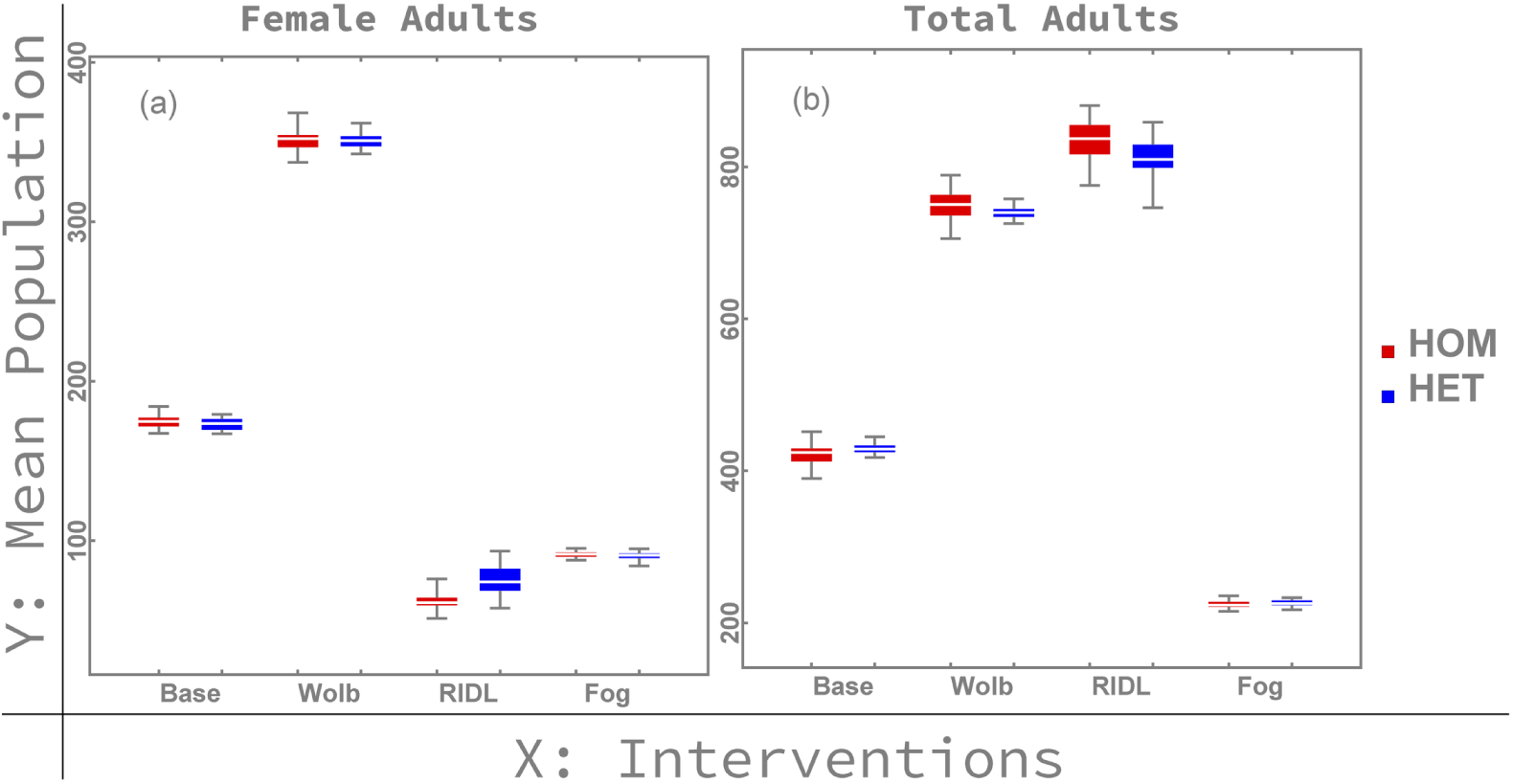
Adult mosquitoes mean population sizes. The area under the curve of the number of adult mosquitoes was calculated for each repetition of each scenario and was divided by the simulated time to obtain these quantities. Figure (a) Shows the number female adults while (b) shows the total adults count. Given this scenarios there was almost no change in the mosquito population sizes of the interventions in their homogeneous setting compared to the heterogeneous one.

Overall, the absence of significant differences in population dynamics due to spatial heterogeneity indicates that difference in mosquito-human vectorial contact networks can be attributed to changes to the spatial arrangement of the environment rather than change in population sizes alone.

### Vectorial-Contact Networks

Given that we have established that there were no significant differences on mosquito population sizes due to spatial distribution changes, we move on to analyse the effect of the spatial layout upon the resulting contact networks. We did find significant differences in the networks between experimental scenarios, so we will divide this section into the analysis of the homogeneous and heterogeneous settings to better highlight the obtained behaviours.

#### Homogeneous Layout

Under this spatial distribution we expected the contact networks to be uniformly distributed as a consequence of each human having the same probability to be bitten. To visually investigate this hypothesis we present the networks’ transition matrices on figures 5a through 5d. To confirm the lack of distinct hosts communities we performed spectral clustering analysis on the networks. No structures on the connections between individuals were found, implying that no bias existed in the way mosquitoes selected their victims. The small-worldness values of these networks also confirm this result as they approach a value of 1 (figure 6c), corresponding to the case where mean path length is equal to the clustering coefficient (which is expected in a uniform network). As a consequence of these results, the degree probability distributions showed a concentrated peak in their of the probability distribution frequencies (PDF) and a sharp transition in the cumulative distribution frequencies (CDF). These outcomes are represented by the solid lines in figure 7. Epidemiologically speaking, this is relevant because these peaks raise the herd immunity threshold to untenably high levels. Under the homogeneous setting of a fully connected network, quarantine ceases to be an effective method to halt transmission as all individuals are highly connected and a large number of edges would need to be removed to disconnect the network (figures 6a and 6b).

**Fig 5.**
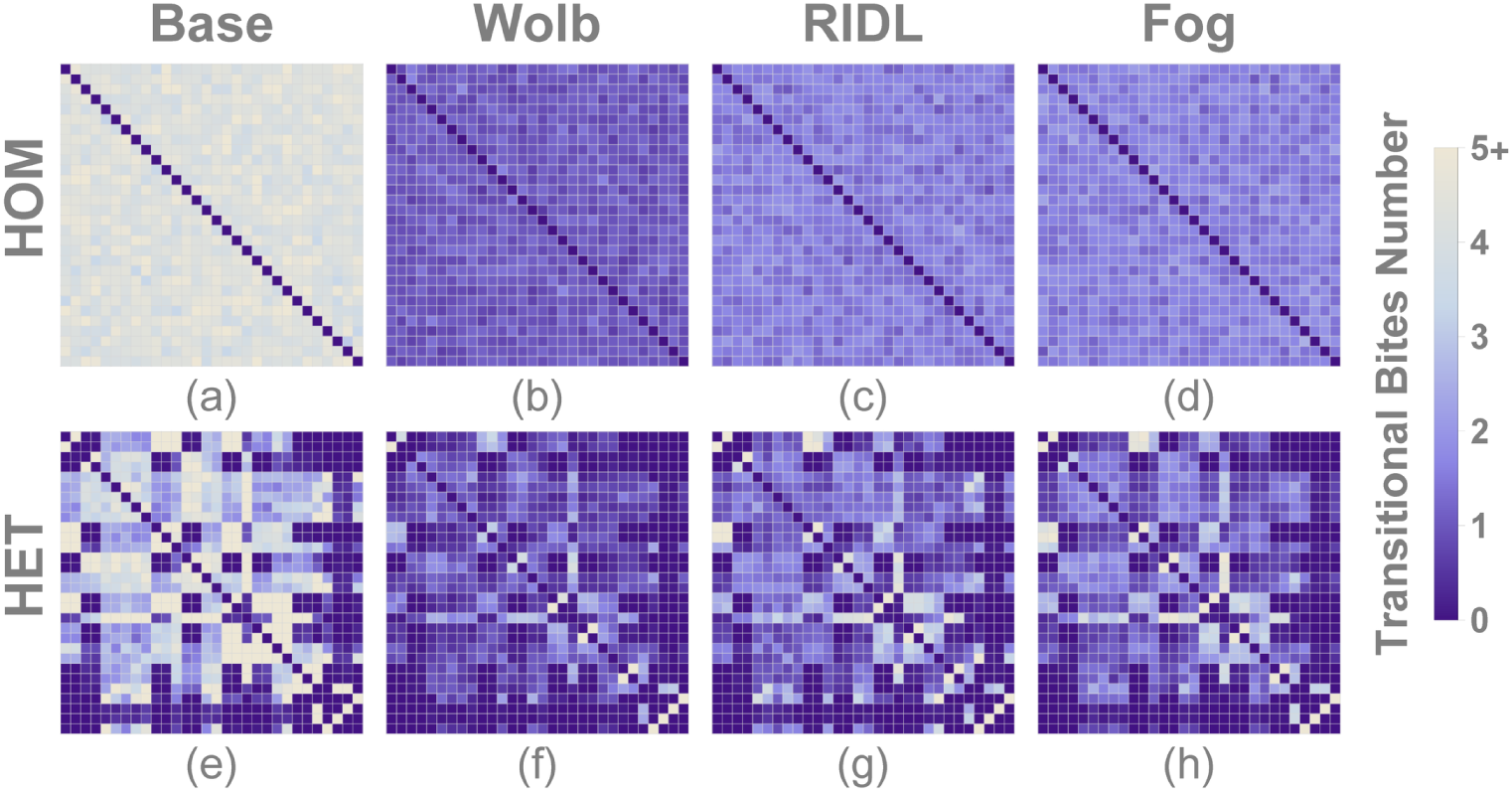
Vectorial contact transition matrices between individuals across different experimental settings. Each row-column intersection is a count of total consecutive bites where the first bite is on the individual indexed by row and the second bite is on the individual indexed by column. In every case the homogeneous setting (top) showed a random pattern while the heterogeneous one (bottom) displayed clear clusters of frequent transitions.

**Fig 6.**
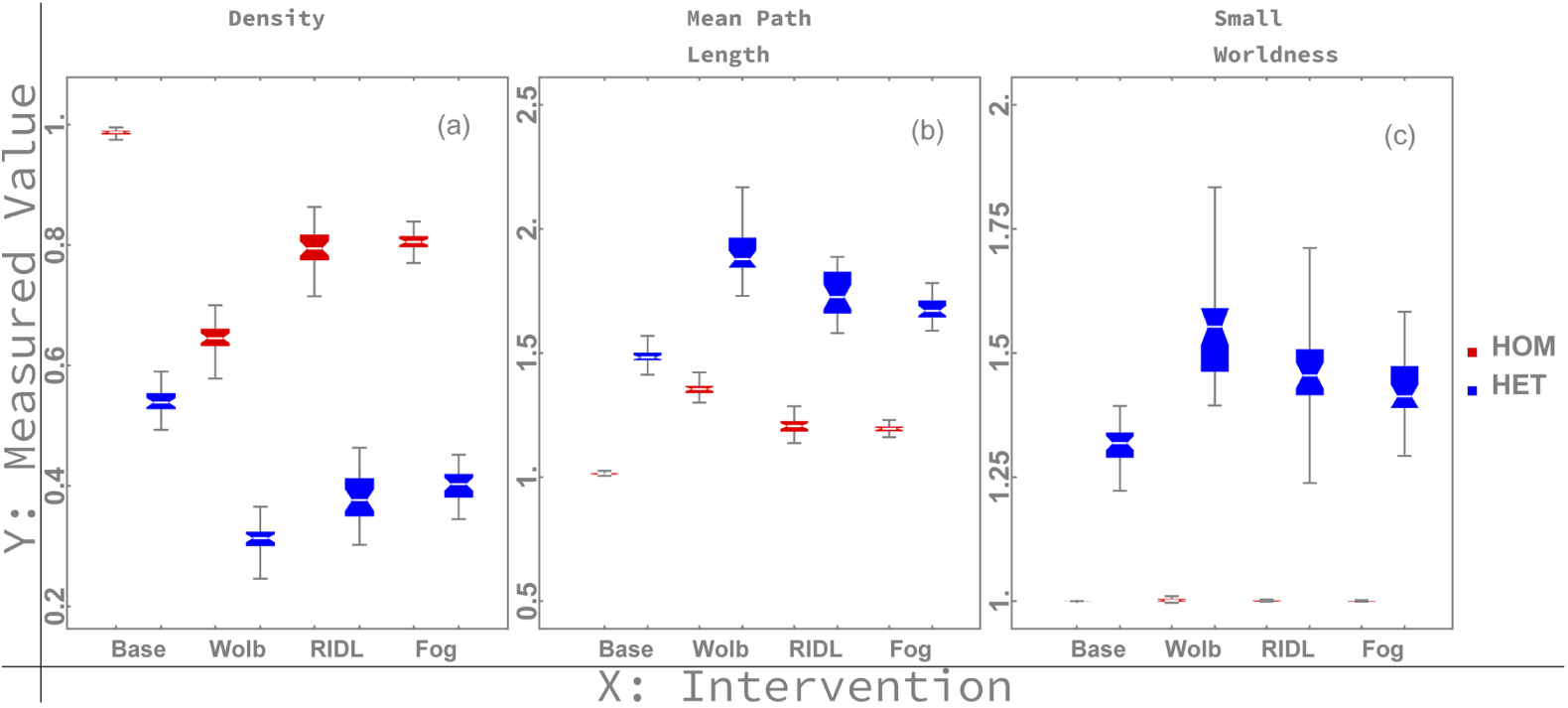
Network measures of spatial scenarios with mosquito-control interventions taking place. Each measure is presented in both their homogeneous (red) and heterogeneous (blue) spatial setting.

**Fig 7.**
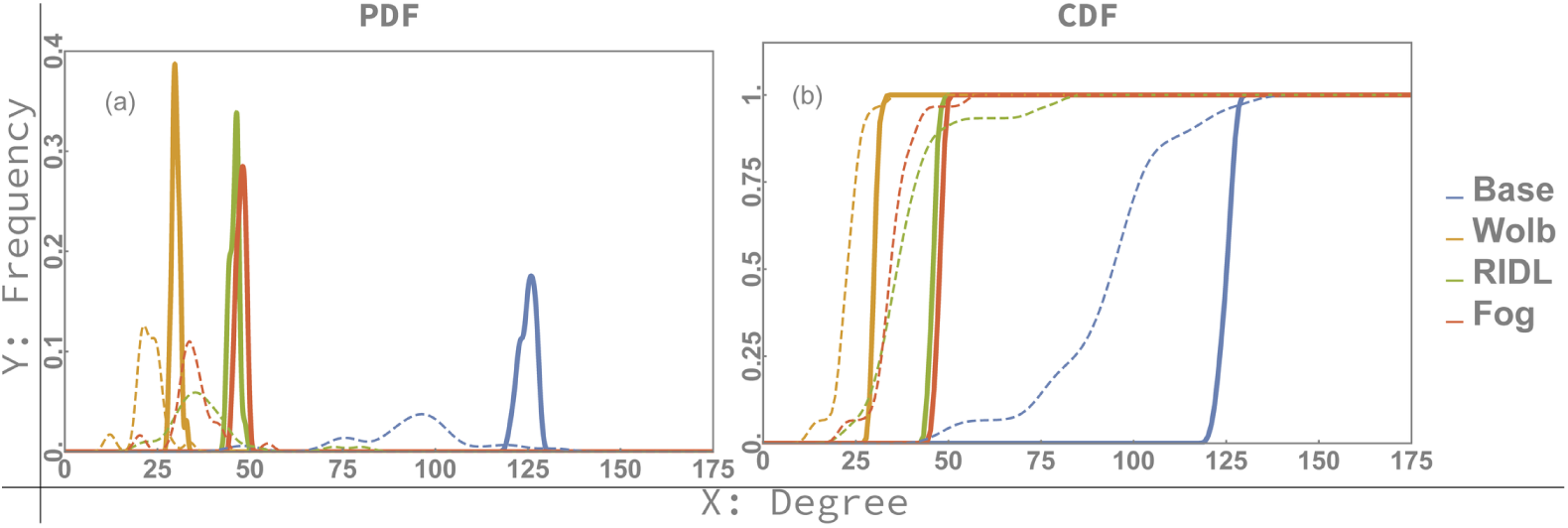
Degree probability distributions of spatial scenarios. Dashed lines represent the heterogeneous scenario while solid lines represent the homogeneous one. PDF’s (a) of homogeneous settings show a more concentrated spike in their transition frequencies which translates into sharper transitions in their CDF’s (b).

#### Heterogeneous Layout

In contrast with the homogeneous settings, these scenarios produced clear patterns in their transition frequencies matrices. The transition matrices shown on figures 5e through 5h clearly show the existence of clusters of individuals (this is a natural effect of *Aedes aegypti* mosquitoes being relatively weak flyers as compared to other mosquito species). We can also see this on figure 8, where some transitions connect individuals more frequently than others (represented by darker lines on the network visualisations). Performing spectral clustering on these networks did find communities that correlate strongly with the spatial distribution of individuals (shown in figure 9). These results allow us to infer that a pathogen would be able to spread with relative ease within these communities, and that targeting the inter-community connections is a better approach to reducing transmission in a population (further demonstrating the idea that human movement plays a major role in dispersing *Aedes*-borne diseases [15, 44]). The close relationship between spatial arrangement of individuals on a landscape and communities embedded in the vectorial network structure hints at the small-world feature to the vectorial contact network. Investigating these network structures should be a priority in larger settings as they are highly relevant to infectious disease epidemiology [39].

**Fig 8.**
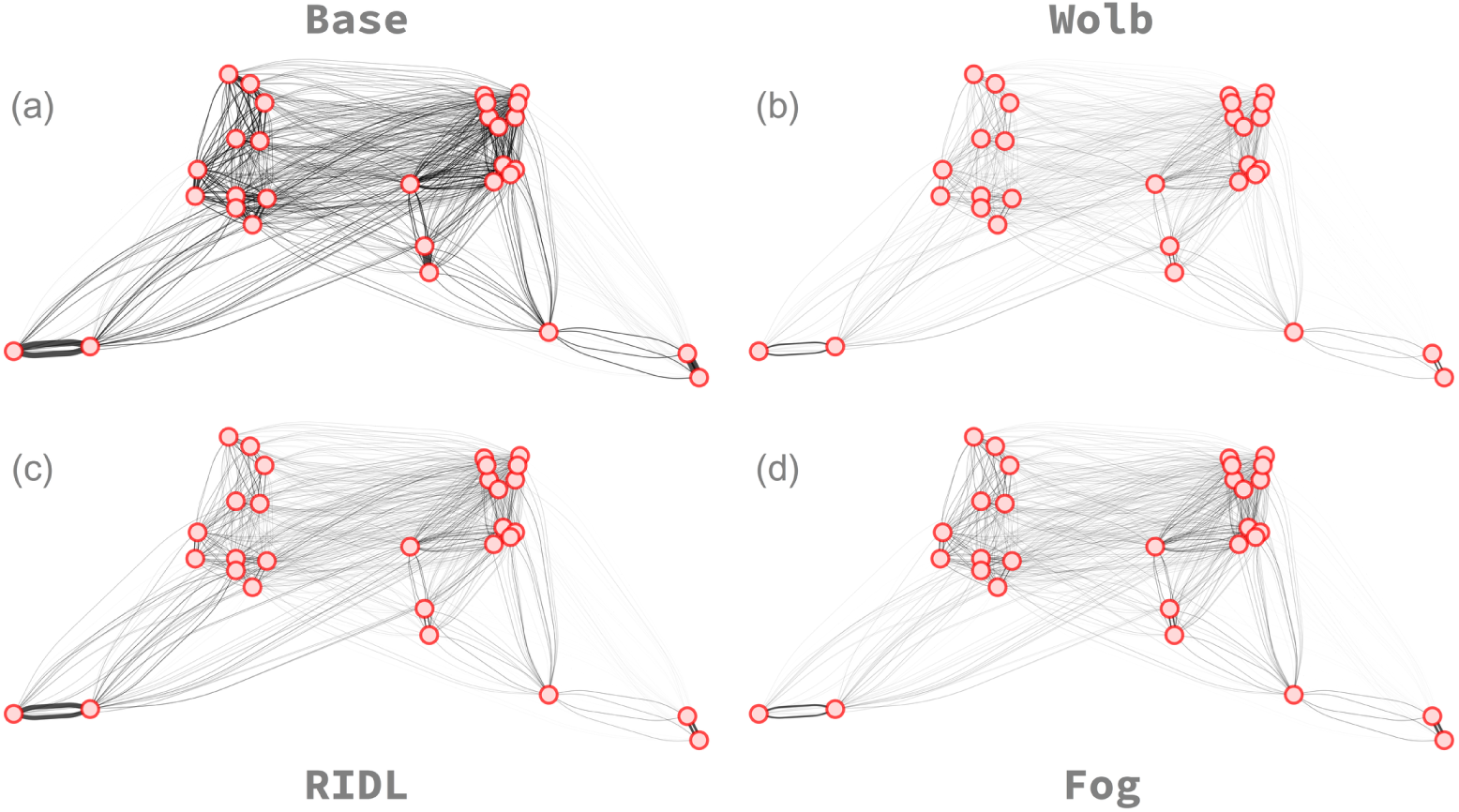
Heterogeneous scenario transition networks. Each node represents a human host and they are spatially distributed according to their location in the simulation. It can be observed that people who spend more time together tend to form stronger links between one another creating tighter clusters of people that live in proximity.

**Fig 9.**
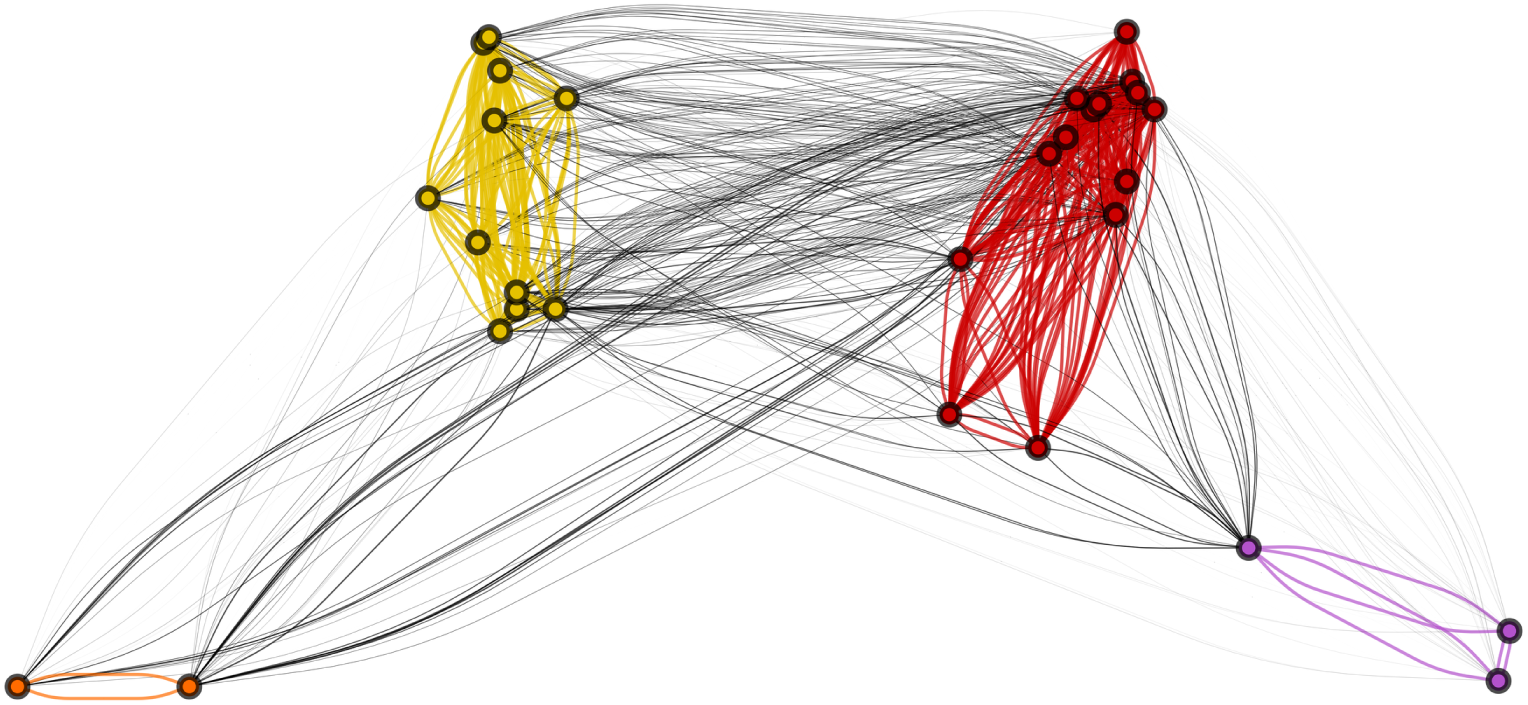
Network communities example. The colour of the nodes represents the community identity as identified by spectral clustering. The homogeneous case contains only one community (omitted) while the heterogeneous one shows that the spatial distribution affects the frequency of transitions and ultimately how the communities are structured. Applying spectral clustering returned the same result on all the interventions’ networks.

Moving on to the degree distributions, the PDF showed a flatter, more platykurtic shape (and the CDF a leaner slope); in the heterogeneous settings than the homogeneous ones (dashed lines in figure 7). Lower network densities and lower connectivities are also related to this effect (figures 6a and 6b), which are relevant because the networks’ capabilities to transmit diseases less robust. Furthermore, all of the interventions’ distribution curves shifted towards the left mainly as a result of the eneral decrease of the number of transitional bites between individuals. This result is confirmed by the networks densities calculations shown in figure 6a in which in every setting has a lower number of transitions on the heterogeneous spatial setting (due to mosquitoes taking more time finding hosts and producing a higher number of self-loops which were discarded according to our proposed methodology).

### Interventions Effects

As the last part of our results description, we will briefly describe the differences on the effects of the control interventions on both the population sizes and networks structures. It should be noted, though, that this is not intended to be a thorough description of the differences between the effects of mosquito control interventions. More variables would be needed to do an analysis of such nature (such as: release distributions, efficacy uncertainties, weather effects, etcetera), but we can make some general assertions of what to expect in a broad sense with the experiments we performed as part of this work. As in previous sections, we will first describe effects on population dynamics and then move on to the networks analyses.

#### Population Dynamics

We can observe on figure 3, that fogging rapidly decreased population sizes from the moment of first application, but that this decrease quickly stabilised to a new equilibrium point after a few treatment repetitions. RIDL releases, on the other hand, showed slower initial decrease of female population size but achieved near total population suppression. In the case of *Wolbachia*, both male and female populations grew, as mixed releases are required due to cytoplasmic incompatibility’s transmission mechanism (it is important to note though, that with *Wolbachia* the goal is not so much to eliminate the mosquitoes population as it is to achieve fixation of the pathogen; so its effects are better described by examining its effect on the vectorial contact networks).s

As a consequence of this analysis we can say that, in the face of sudden epidemic episodes, fogging might be a viable alternative towards quickly reducing the opportunities for the pathogen to spread. To combat endemic pathogen transmission, however, we would want to shift towards either the eradication of mosquitoes through RIDL or the fixation of *Wolbachia* to disrupt pathogen transmission. This can be done after reducing population sizes through more traditional approaches such as source reduction or fogging (which falls in line with how these two interventions are usually applied on the field or designed to work [13, 14]).

#### Vectorial-Contact Networks

Networks were sparser in all the heterogeneous scenarios with respect to the homogeneous ones, but *Wolbachia* produced the largest effect overall on lowering their densities and degrees (figures 6a and 7). Despite this, it is interesting to point out the behaviour of its small-worldness value. Although *Wolbachia* showed good qualities in disrupting the transmission network (along with higher mean path length values, as shown in figure 8b), it also showed the highest small-world coefficient; which would imply that the mean path length of the network would scale as the logarithm of its number of vertices, keeping the persons epidemiologically “close” to each other even while human population grew in number (given that they scale in similar spatial and behavioural patterns).

In terms of degree probability distributions, we can see the emergence of several interesting behaviours. The baseline scenario presented more heterogeneity in the transitional biting behaviour between landscapes (a more flatter shape on the distribution on figure 7a). This is probably due to the fact that more mosquitoes were able to survive and create some sporadic long distance transitions between humans (effects which are dampened in the cases where the interventions are applied). RIDL managed to reduce its PDF peak to a lower value than fogging and *Wolbachia*; while the latter was the one with the largest change between spatial settings. In general terms, a more heterogeneous the number of bites would mean that the bites are concentrated amongst certain individuals in the network, individuals which could be targeted to reduce diseases’ spread (by using it’s centrality as a proxy measure of this “importance” in the epidemiological structure). Taking this into account RIDL could be the intervention with greater effect, although more repetitions would probably be required to make the distribution frequencies converge into more stable shapes for definite conclusions to be made.

## Discussion

After comparing mosquito population dynamics and vectorial-contact networks it is evident that while overall population counts are useful to produce rough estimates of expected level of potential disease transmission they are less useful to examine how a pathogen can spread through a host population in a spatially heterogeneous scenario. This insufficiency of mean population counts to provide useful information in the face of spatial and other heterogeneities becomes even more evident when considering the effect of vector control interventions. Vectorial contact networks, on the other hand, are able to give a precise mathematical description of how an *Aedes*-borne pathogen might spread in a spatially distributed host population. However, calculation of these vectorial-contact networks in the field is operationally unfeasible, motivating our proposal to use detailed agent-based simulations to further our understanding of how epidemic processes may occur on real landscapes.

The vectorial-contact transition matrices derived from our simulations provide a precise mathematical description of how hosts are epidemiologically connected through vector contact. These matrices therefore give detailed individual level form of classic transmission metrics such as *R*0 and vectorial capacity [45]. While under certain limiting circumstances transmission dynamics could be well described by mean-field approximations based on systems of ordinary differential equations, finite population sizes, heterogeneous biting, and spatial aggregation patterns found in real transmission settings might invalidate these mean-field assumptions. While sophisticated mathematical techniques such as spatial moment-equations could be used to incorporate spatial effects into a deterministic model of transmission, assumptions must still be made in order to keep the models analytically tractable. Especially in settings characterised by heterogeneities of host behaviour and spatial distribution, as well as small population sizes, commonly encountered in residual transmission scenarios, it is paramount to capture emergent properties of the transmission dynamics which highly depend on these peculiarities of the setting. In these cases agent-based simulation provides an effective means by which transmission dynamics on real landscapes can be easily simulated and analysed.

Results from our spatially-explicit agent-based simulations strongly indicate that heterogeneous spatial distribution of hosts and mosquito breeding sites greatly impacts how a pathogen may invade a human population when mediated by *Aedes* mosquitoes. These differences in the epidemiological relations between individuals is clear from figures 5 and 7 where the inclusion of spatial heterogeneity produced drastically different epidemiological settings. While in all cases the spatially homogeneous scenario produced the worst-case scenario across all interventions (figures 6 and 7) this simulated scenario is of marginal use in planning vector or host based interventions in the field. In many cases interventions aimed to mitigate the worst-case scenario may be much more costly and have much less impact per dollar spent than a targeted intervention informed through analysis of realistic spatially heterogeneous simulations. Furthermore, analyses of simulations under the assumption of spatial homogeneity lose relevance when considering scenarios of low-prevalence and residual transmission. The importance of spatial distribution to vectorial-contact may be observed in figure 9. The network structure can be observed to be characterised by several dense clusters of individuals that correlated strongly with the spatial distribution of hosts and breeding sites. This partitioning of the network into tightly connected clusters suggests that vector-borne pathogens can spread efficiently within clusters. These clusters may support residual pathogen transmission and provide a reservoir for disease even when other clusters or areas of the terrain are successfully targeted by transmission control campaigns. This is most relevant when considering elimination scenarios because these pockets of disease provide the pathogen a means of persistence and possible re-emergence even if inter-cluster transition probability is low (due to the small-world nature of the contact networks in the heterogeneous setting as shown in figure 6c).

With respect to comparison of different vector control interventions applied to the spatially heterogeneous scenario there was no evidence of substantial differences that were only attributable to spatial effects, although one notable difference was the small-worldness of the vectorial-contact networks, which seemed to present different behaviour on each intervention, shown in figure 6c. Most of the variance in calculated measures when compared between interventions can be solely attributed to reduction in population size (this can be seen in figure 6; measures closely follow the distributions in figure 7 which themselves strongly depend on overall vector population density). This is to be expected according to our experimental design where vector control interventions were applied uniformly to the simulated landscape. We simulated the somewhat unrealistic assumption of uniform application of interventions in order to compare their effects on population density and network measures without potential confounding from spatial distribution of the interventions themselves. In future research we plan to preform a more thorough analysis of each intervention including realistic spatial applications (targeting mosquito mating swarms, hotspot releases, rearing releases, et cetera). Analysing the interventions under realistic operational constraints should provide a better picture of how vector control interventions can be targeted to take advantage of spatial heterogeneity in host distribution with respect to specific properties of each intervention to maximise their impact on fragmenting vectorial-contact networks.

We performed the aforementioned analysis to demonstrate the importance of acknowledging spatial distribution of hosts and breeding sites when planning vector-control interventions for *Aedes*-borne pathogens. We note however, that much work is still required to produce definite conclusions of how disease spread is affected by spatial heterogeneities. In particular, we plan on extending our model to accommodate larger human population sizes, more realistic mosquito-control releases, data-informed human movement and pathogen models; to be able to make location-specific analyses on how to control epidemic processes efficiently.

## Conclusions

Understanding the effects of spatial heterogeneity in mosquito-borne diseases is a difficult task, but with the use of agent-based models and network theory we have shown that it has a significant effect on how humans connect to each other through *Aedes aegypti* mosquito bites both in absence and in presence of three different mosquito-control interventions. This highlights not only the fact that spatial heterogeneity is an extremely important element of the transmission of mosquito-borne diseases, but also the need of new tools to further our understanding of the implications and effects it has on epidemic processes and vector-control interventions.

These initial conclusions are meant to serve as a guide for future research, as much work is still needed to get a bigger picture of how these heterogeneous contacts dynamics emerge from human-mosquito interactions; and how to take advantage of them to limit diseases spread. In particular, we want to simulate larger geographical regions with more realistic behaviours both in human behaviour and in weather patterns, to have a more robust model of the networks that result as a consequence of their interactions with mosquitoes. We are also planning on making a more thorough analysis of how spatial heterogeneity in the application/release of vector-control interventions affects the contact-networks. All of these analyses would help us move towards the efficient use of the limited resources dedicated to the eradication of often neglected tropical diseases transmitted by *Aedes aegypti* mosquitoes.

## Acknowledgments

This work was supported by UC-MEXUS grant. Award number: CN-15-47. Any opinions, findings and conclusions or recommendations expressed in this publication are those of the authors and do not necessarily reflect the views of the sponsoring agencies. Thanks to Julio G. Arriaga, Benjam´ın Vald´es and Victor Ferman for their kind comments and help in the writing of this paper.

